# Variation in echolocation call emission of Neotropical insect-eating bats in response to shifting ambient temperatures

**DOI:** 10.1101/2023.12.20.572652

**Authors:** Paula Iturralde Pólit, Marcelo Araya-Salas, Holger R. Goerlitz, Gloriana Chaverri

**Author notes:** these authors are joint senior authors (Holger R. Goerlitz, Gloriana Chaverri).

## Abstract

The sensory systems of animals are essential for them to respond to environmental cues and signals. However, their functionality might be altered by climate change. Most bats, for example, rely on acoustic signal emission for acquiring food, but their high-frequency echolocation calls are strongly attenuated in the air. Attenuation in air changes with changing weather conditions, which can lead to shifts in echo-based prey detection distance. However, bats adjust call parameters to the task and environment, and this behavioural plasticity may also help them to counteract potential increases in sound attenuation to keep echo detectability constant. We explored this ability in a community of insectivorous bats in a montane forest of Costa Rica. We recorded bat echolocation calls in response to experimentally increased temperatures, simulating intermediate and arguably realistic projected climate change scenarios. We calculated atmospheric attenuation and detection distance for each temperature and echolocation call. We found some changes in source level and call duration, yet not in peak frequency, and responses were not consistent across species with increasing atmospheric attenuation. This might be explained by several non-mutually exclusive reasons, including that the experimental increase in temperature and change of atmospheric attenuation were not sufficient to affect close-range prey detection. Ultimately, this study contributes to our understanding of sensory system adaptation under the pressure imposed by climate change.

**SUMMARY STATEMENT:** Studying adjustments in bats’ call parameters can reveal responses to the pressure imposed by climate change.

## INTRODUCTION

Climate change is affecting biodiversity in various ways; however, most research has been mainly focused on assessing shifts in potential distributions of local and invasive species (Anderson, 2013; Crespo□Pérez et al., 2015; Gallardo et al., 2017; Iwamura et al., 2020; Pecl et al., 2017; Peterson et al., 2018), species phenology (Parmesan, 2006), and species vulnerability (Foden et al., 2013; Williams et al., 2008). In addition, available data are geographically biased (Martin et al., 2012; Meyer et al., 2015), precluding us from accurately predicting its effect more broadly. Many climate change studies reference global response patterns, yet latitudes below 30° remain underrepresented (Feeley et al., 2017) despite being the areas with the greatest biodiversity (Barlow et al., 2018). Furthermore, those studies often forecast shifts based on correlations of available information, but they rarely consider the mechanisms that shape species’ responses to increasingly warming conditions (Urban et al., 2016). This can only be assessed with experimental studies, which are crucial in determining the ability of species to respond to climate change (Razgour et al., 2019; Willis et al., 2015). For example, understanding potential changes in a species’ perceptual abilities under changing conditions provides insights into its ability to overcome environmental shifts, but these studies are still scarce in tropical regions (Beever et al., 2017).

Sensory ecology studies how organisms acquire and respond to environmental cues and signals. Animals rely on this information for courtship, individual recognition, orientation, and prey detection, among others (Stevens, 2013). Unfortunately, the functionality of animal sensory systems may be altered by several factors, for example by noise pollution both in terrestrial (Dominoni et al., 2020; Tuomainen & Candolin, 2011) and aquatic environments (Johnson et al., 2009; Kelley et al., 2018). Most recently, and while data are still scarce, several studies suggest that anthropogenic climate change may be an additional factor that affects animals’ sensory perception. For example, in aquatic environments, higher CO_2_ concentrations and ocean acidification can impair olfactory sensitivity in fish, reduce the efficiency of visual cues that indicate predator presence, and disrupt auditory-guided behaviours (Draper & Weissburg, 2019; Kelley et al., 2018; Porteus et al., 2018; Rivest et al., 2019). Other effects of climate change, such as changing temperatures, could affect terrestrial animals. For instance, lizards, moths, and flies could suffer a reduction in the effectiveness of chemical sexual signals, which are essential for mate choice (Groot & Zizzari, 2019; Martín & López, 2013).

Sound is a central stimulus for many animals to acquire environmental information (Dusenbery, 2001). For example, echolocation is a sensory mechanism by which animals produce high-frequency vocal signals and listen to the returning echoes to perceive their surroundings. Echolocating animals, like bats, use this mechanism for spatial orientation and often to detect, localize, and intercept prey (Schnitzler & Kalko, 2001), and very flexibly adjust their signals to their current behavioural task and to habitat conditions (Neuweiler, 1989; Schnitzler et al., 2003). The emitted calls of echolocating bats may also be influenced by weather conditions, as temperature and humidity play a significant role in the atmospheric attenuation of the high-frequency calls commonly used by these mammals (Goerlitz, 2018). In general, adapting calls to different environmental situations may reduce excess attenuation through scattering (Römer, 2001), optimize sound propagation, and increase foraging success (Luo et al., 2014; Obrist, 1995; Surlykke & Kalko, 2008).

Changing weather conditions, either originating from short-term spatio-temporal variation or from long-term climate change, affect the volume of space over which bats can detect prey (Luo et al., 2014). Hence, bats may rely on vocal plasticity to compensate for this effect. While some species show long-term acoustic signal divergence associated with adaptation of sensory systems to local environmental conditions (geographical variation in average weather parameters, Chen et al., 2009; Maluleke et al., 2017; Mutumi et al., 2016), individuals can also plasticly adjust their signals in response to seasonal and daily weather fluctuations (Chaverri & Quirós, 2017; Snell-Rood, 2012). These findings show that bats have control over their acoustic signals, making them a good study system to investigate adjustments in vocal production in response to the predicted shifts in weather due to climate change.

To date, studies focusing on sensory responses of bats to fluctuations in atmospheric conditions on a short temporal scale are scarce. Some studies have assessed effects on detection distance (de Framond, Reininger, et al., 2023) and adjustments in echolocation calls in a temperate region (Snell-Rood, 2012). To our knowledge, only one study has evaluated the association between weather conditions and acoustic parameters of calls in echolocating Neotropical bats (Chaverri & Quirós, 2017). Neotropical species, particularly those in higher elevations, have received little attention despite living in a region predicted to experience substantial changes in weather (Boehmer, 2011; Still et al., 1999) and significantly higher maximum temperatures (Enquist, 2002). In addition, the more constant weather conditions in the tropics compared to temperate regions contribute to narrower thermal tolerances, smaller distribution ranges, higher species turnover along altitudinal gradients (Ghalambor et al., 2006), and higher levels of endemism (Chaverri et al., 2016). Altogether, these factors could render tropical bat species more sensitive and vulnerable to climate change, posing a potential threat to their long-term survival.

Here, we aim to gather empirical data on the acoustic responses of a community of Neotropical insectivorous bats to changes in the abiotic environmental conditions known to affect sound transmission. We hypothesized that echolocating bats modify their call parameters in response to changing atmospheric conditions, which affect sound attenuation and thus maximum detection distance. In response to increasing atmospheric attenuation, bats might either decrease call frequency, increase emitted call level, increase call duration, or any combination thereof, in such a way to maintain maximum detection distance (Luo et al., 2014; Snell-Rood, 2012). We simulated increasing temperatures in line with climate change scenarios and recorded the bats’ echolocation calls, analysed their call parameters, and calculated the resulting atmospheric attenuation and maximum detection distance, to assess the influence of atmospheric attenuation on call emission. Because environmental conditions are often highly variable, assessing vocal flexibility in bats at short to medium temporal scales will help predict species sensitivity and resilience to some of the abrupt shifts in weather conditions caused by climate change (Alley et al., 2003).

## MATERIALS AND METHODS

### Field site and species

We collected data at Las Cruces Research Station located in Southern Costa Rica at Coto-Brus county, close to the boundary of the largest protected area in the country (Parque Internacional La Amistad). The site includes tropical pre-montane and lower montane forests with frequent presence of clouds (Enquist, 2002). Altitude ranges from 1200 to 1800 m above sea level, ambient temperature between 15°C at night and 28°C during the day, and relative humidity fluctuates between 60% and 100% throughout the year. These ecosystems are crucial for biodiversity conservation due to high species diversity and endemism (Cadena et al., 2012), where bats are not the exception (Chaverri et al., 2016; Pineda-Lizano & Chaverri, 2022). However, it has been predicted that the Neotropical region will suffer significant climate change-induced shifts in weather conditions of up to 4°C with slight differences between the wet and dry seasons (Ambarish & Karmalkar et al., 2011) and that highland sites may suffer some of the most significant effects on biodiversity (Karmalkar et al., 2008; Mata-Guel et al., 2023).

We focused on insectivorous bats from the genera *Myotis* and *Eptesicus* (family Vespertilionidae). These bats often fly in the understory (Kalko et al., 2008) and are mainly edge-space foragers, yet also move between different types of habitats when searching for prey, thereby adapting their echolocation calls to different foraging situations (Denzinger & Schnitzler, 2013).

We captured bats with mist-nets from 17:45 (before dusk at the study site), when vespertilionid bats started their activities, until 21:30-22:00, when the first bout of bat activity dropped. We aged, sexed, and taxonomically identified all individuals, then fed them with mealworms (larvae of *Tenebrio molitor*), provided water *ad libitum,* and kept them individually in cloth bags until the start of the experiments (around 22:30 – 23:00).

Taxonomic identification of species in the field can be difficult, especially when species are cryptic, as is the case with some *Myotis* species (York et al., 2019). Since genetic identification was not possible, and following the Field Key to the Bats of Costa Rica and Nicaragua (York et al., 2019), we identified *M. oxyotus* and *M. riparius* based on the arrangement of the first two premolars. For *M. pilosatibialis*, we verified that the uropatagium and legs had fur at least to the knee. Identification of *Myotis nigricans* and *M. elegans* was more problematic, and we collected few individuals of these species because they are more common in the lowlands. To avoid misidentification, we grouped these individuals (herein *Myotis nigricans/elegans*) for further analyses.

### Experimental setup

We recorded the echolocation calls of individual free-flying bats in an outdoor flight cage (6 x 2.5 x 2.5 m^3^) with a four-microphone array (Fig. 1). The cage walls were made of a synthetic cloth, reducing the exchange of air between the cage and the environment. To reduce reverberations, the cage walls and ground were covered with sound-absorbing fabric, with additional sound-absorbing foam on the ground.

**Figure 1:**
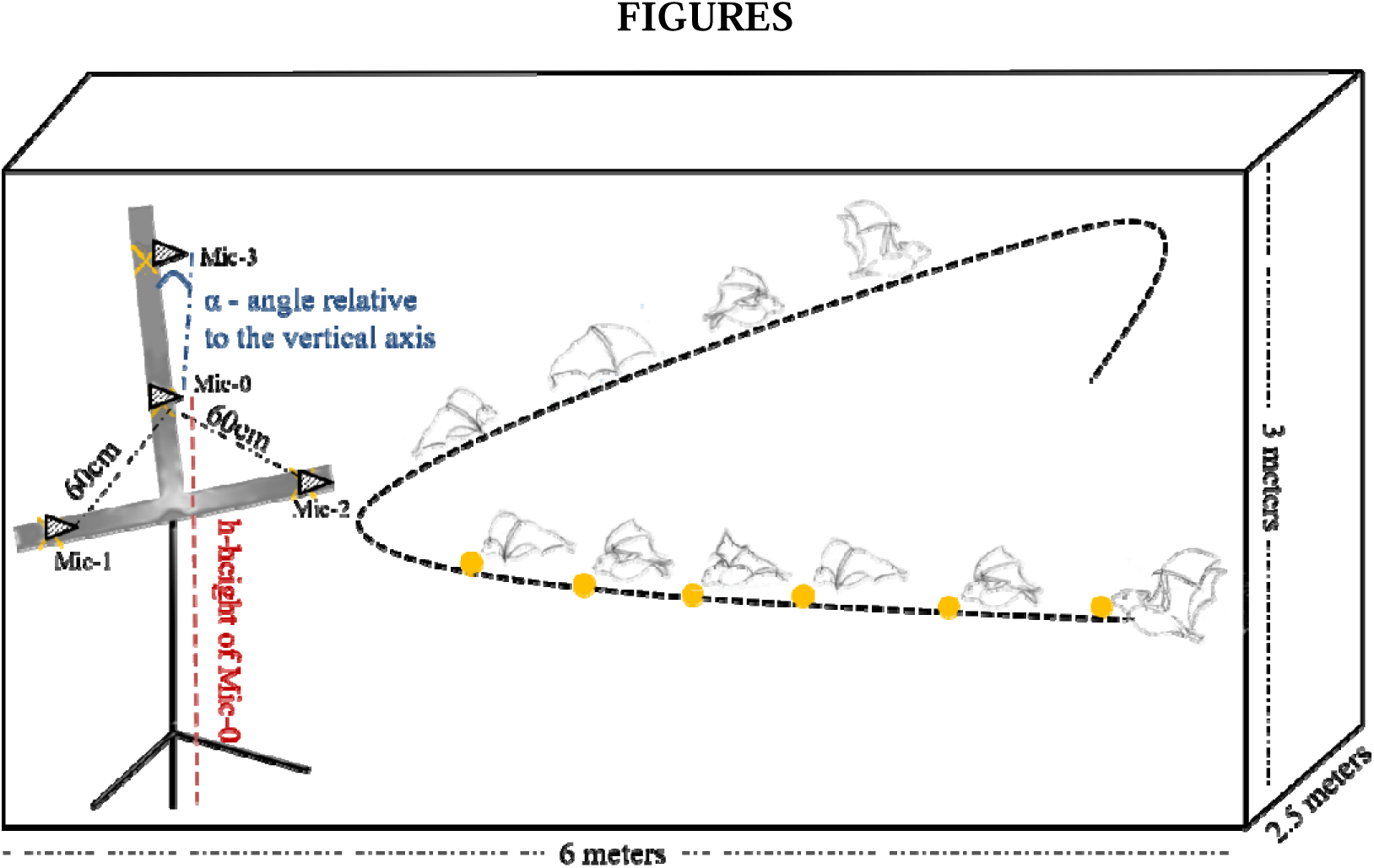
Experimental setup. Bats were flying freely and individually inside a flight cage. Their calls were recorded with a four-microphone array, which was placed on one side of the flight cage. The microphones were arranged in a star shape within the T-shaped structure.

We positioned the microphone array centrally at one side of the flight room to record the echolocation calls of the bats. The array consisted of four omnidirectional electret ultrasound microphones (Knowles FG-O, Avisoft Bioacoustics, Glienicke/Nordbahn, Germany) arranged in a symmetrical star-shape and mounted on a T-shaped metal structure covered with sound-absorbing foam. The three outer microphones had a distance of 60 cm to the central microphone. Microphone signals were recorded with an Avisoft Ultra Sound Gate 416 H and Avisoft Recorder software to a four-channel WAV-file at 500 kHz sampling rate and 16-bit amplitude resolution, and maximum gain without clipping the calls.

We calibrated the frequency response and the directionality of the central microphone, which was used for call analysis, before the experiment. We played pure tones with constant frequency from 5 to 95 kHz in steps of 5 kHz from a loudspeaker (Vifa; Avisoft Bioacoustics) and recorded them with a measuring microphone with flat frequency response (G.R.A.S. Sound and Vibration A/S, Holte, Denmark) placed at 50 cm distance. We recorded the same pure tones with the central microphone from directions from 0 to 90° in 5° steps. By comparing recordings on the measuring microphone and the central microphone, we obtained the frequency response and directionality of the central microphone, which we used to correct each recorded call to obtain the call as received at the microphone, which was further corrected to obtain the call as emitted by the bat (see “Estimating call parameters” below).

### Recording procedure

To investigate if and how increasing temperature affects call parameters, we recorded the echolocation calls of individual free-flying bats under three different temperature conditions. The least and most extreme climate change projections (Pachauri et al., 2014) predict an increase in ambient temperature of 1°C (RCP2.6) and 4.8°C (RCP8.6), respectively. We aimed to simulate the intermediate and arguably more realistic scenarios RCP4.5 and RCP 6.0, which predict temperature increases of 2°C and 4°C.

We tested each bat in one night by presenting three different conditions (current ambient temperature Ta, Ta + 2°C, and Ta + 4°C). First, we recorded each bat under unmodified ambient atmospheric conditions (Ta), in which the temperature and humidity of the flight cage were similar to external conditions. For the two subsequent trials, we increased the temperature in the flight cage by 2°C and 4°C, respectively, using two electric heaters placed in the middle of the flight cage. Once we reached the target temperature, we removed the heaters from the cage to reduce reverberations.

Temperature and relative humidity in the flight cage were continuously recorded with a weather logger (Kestrel 4000, pocket weather tracker; KestrelMeters, Boothwyn, Pennsylvania, USA). Each bat was tested under all three conditions, and in the same sequence. If more than one bat was captured in one night, we first tested all of them individually under Ta, and then under the sequentially increased temperatures.

For each of the three trials, we released the bat into the flight cage for no longer than five minutes and recorded between five to ten audio files of approximately 10 seconds each. After each trial, we caught the individual with a hand net, fed it with mealworms and provided water *ad libitum*. After the experiments, the bats were released at their capture site.

### Estimating call parameters

We checked all recordings using SASLab Pro (Avisoft Bioacoustics) and estimated acoustic parameters (maximum and minimum call frequency, call duration, and inter-call interval) per species. We used this information to determine an appropriate bandpass filter for every recording session for final call analysis. We used the custom- developed TOAD Suite software package (de Framond, Beleyur, et al., 2023; de Framond, Reininger, et al., 2023; Hügel et al., 2017; Lewanzik & Goerlitz, 2018) for MATLAB (Version R2007b; The Mathworks, Inc., Nattick (MA), USA) to calculate the bats’ spatial position for each emitted signal based on the time-of-arrival differences (TOAD) from the central to the outer microphones of the array (Koblitz, 2018) and the speed of sound for the current air temperature and relative humidity.

We then reconstructed the 3D flight trajectories, and manually selected at least four consecutive calls without overlapping echoes (quality calls) from trajectory segments where the bats were flying towards the microphone array. All selected calls were automatically corrected for atmospheric absorption and spherical spreading on the way from the bat to the microphone and the frequency response and directionality of the microphone, to obtain the call as emitted by the bat at 10 cm from its mouth. We then automatically calculated call duration based on the smoothed Hilbert envelope at -12 dB relative to the envelope’s peak amplitude value; peak frequency (the frequency with the highest amplitude); and apparent source level (aSL) as the root mean square (rms) relative to 20 µPa and at 10 cm to the bat’s mouth (rms dB re 20 µPa @10 cm). Since bat calls are highly directional and not necessarily emitted towards the microphone, the aSL is an underestimation of the real on-axis maximum source level (SL).

We excluded calls with signal-to-noise ratios < 30 dB and those in which the maximum energy was detected in the second harmonic. This resulted in a dataset of 5,104 calls of five species (groups): *Myotis nigricans/elegans* group: 735 calls; *M. pilosatibialis*: 2,567; *M. riparius*: 978; *M. oxyotus*: 331, *E. brasiliensis*: 493.

To approximate the real SL from aSL, we only kept calls above the 90^th^ percentile of the aSL within one experimental trial of one bat. This resulted in a total of 1.002 calls for final analysis (Table 1). The most abundant species in our sample was *Myotis pilosatibialis* with 31 individuals, from which we also recorded the highest number of calls in the final dataset (494).

**Table 1:**
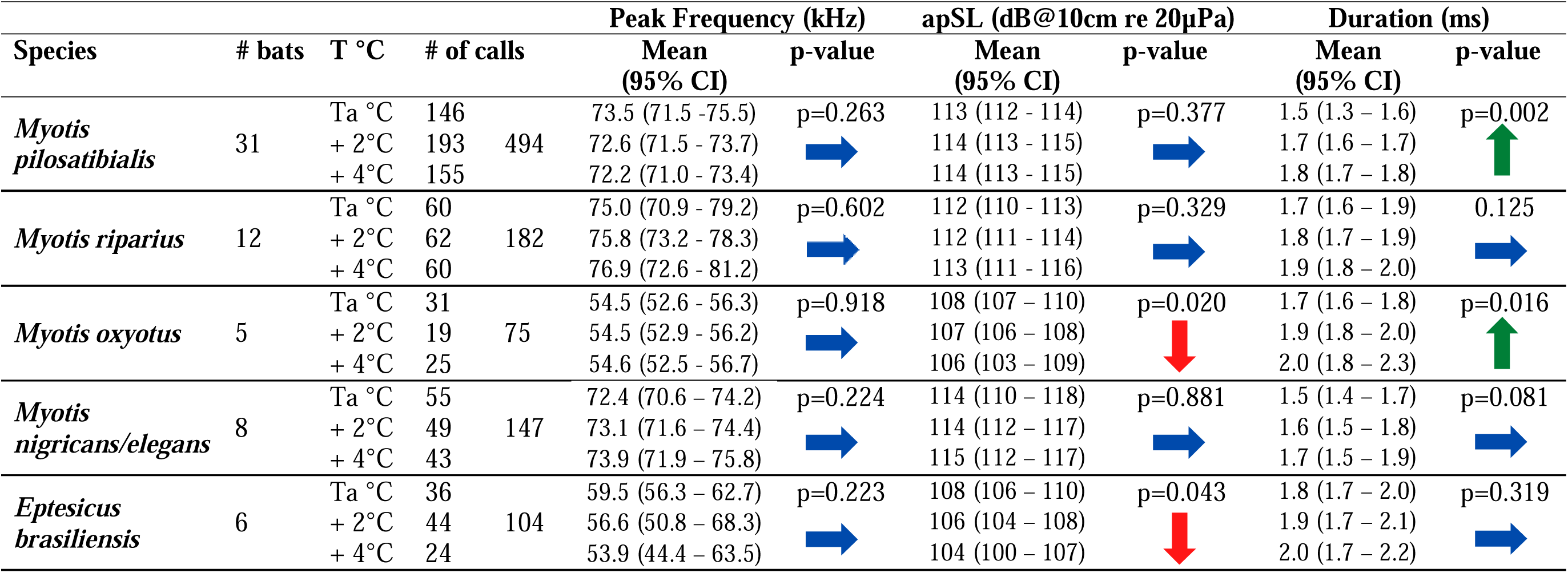
The number of bat individuals and calls per experimental temperature per species (Ta: ambient temperature). We show call parameters per species and temperature condition with the statistical results (p-value) of the random slope models and the direction of change represented with arrows (no change = blue, negative = red, positive = green) of the call parameter with increasing temperature.

### Statistical analyses

#### The effect of temperature on echolocation calls

All statistical analyses were conducted using R version 4.0.5. To determine whether bats adapt call parameters to increased ambient temperatures, we compared two linear mixed-effects models using the R-package lme4, version 1.1-28 (Bates et al., 2020). We calculated random intercept models and random slope models per species as follows:

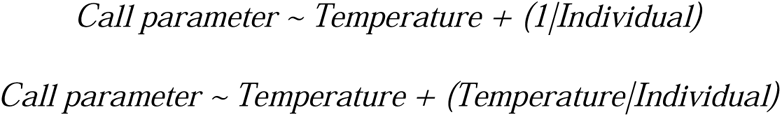

Random intercept models assume that all individuals respond similarly (Bates et al., 2014). However, we found high inter-individual variation in call parameters in all species (figures S1, S2, and S3), suggesting that individuals could respond differently, which is better captured by random slope models. Since the random slope model fitted the data better for three out of five species (Likelihood Ratio Test p<0.05; AIC-difference > 2), we chose them for all species.

#### The effect of temperature on atmospheric attenuation and detection distance

To investigate whether bats adjust call parameters to weather conditions to maintain detection distance, we calculated atmospheric attenuation (AA) of sound and detection distance (DD) for prey based on weather conditions and call parameters. AA describes how much the level of a sound is weakened per distance, expressed in decibels per meter (dB/m), and depends (in decreasing order) on call frequency, ambient temperature, relative humidity and atmospheric pressure (Goerlitz, 2018). DD is the distance over which a bat can detect an object, for example a prey item. According to the sonar equation (Møhl, 1988), DD depends on the emitted apparent source level (aSL), AA, the sound reflectivity of the object (target strength, TS) and the bat’s hearing threshold. Therefore, AA is a function of call peak frequency, temperature and relative humidity; and DD in turn is a function of AA, aSL, prey target strength, and hearing threshold (sonar equation; Møhl, 1988).

Target strength is strongly influenced by prey size (de Framond, Reininger, et al., 2023), its orientation towards the bat (Sümer et al., 2009; Waters et al., 1995), and surface properties (Neil et al., 2020a; Simon et al., 2023). In general, neotropical vespertilionid species prefer soft prey like nocturnal lepidoptera (moths) from a wide range of sizes (Aguirre et al., 2003; Ingala et al., 2021). To account for this variation, we set TS (Møhl, 1988; Surlykke et al., 1999; Surlykke & Kalko, 2008; Waters et al., 1995). Note that these are just approximate values, given the various factors influencing TS (Neil et al., 2020a, 2020b). We set the hearing threshold at 20 dB SPL to account for noise and behavioural reaction thresholds (Boonman et al., 2013).

To separate the effect of changing temperature and changing call parameters, we compared an assumed *non-responsive* bat that uses non-adjusted echolocation call parameters with a *responsive* bat that uses the actual call parameters recorded for different experimental temperatures. We first calculated, per individual, the mean values of peak frequency, aSL and duration at each experimental temperature. For the *non-responsive* bat, we only used the mean call parameters at ambient temperature to calculate AA and DD; thus, we assume these bats do not adjust call parameters with increasing temperatures. For the *responsive* bat, we used the actual call parameters per temperature; thus, we assume call adjustments as temperatures increase. To include the effect of changing call duration on DD, we lowered the *responsive* bat’s hearing threshold by 6 dB for every doubling of duration (Luo et al., 2015, for short calls < ∼2 ms):

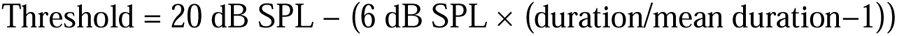

By comparing AA between *responsive* and *non-responsive* bats, we tested whether bats counteracted a potential temperature-induced increase in AA by lowering call frequency. Also, by comparing DD between *responsive* and *non-responsive* bats, we tested whether the bats counteracted a potential temperature-induced decrease in DD by adjusting call parameters, i.e., if they maintained prey detection distance when facing warming conditions.

Using the AA and DD data for the *non-responsive* and *responsive* bat, we calculated an interaction model to determine whether changes in AA and DD over temperature differed between the *non-responsive* and *responsive* bat:

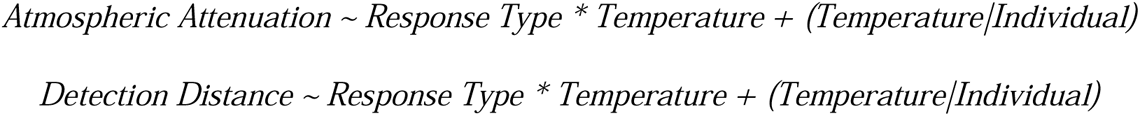

These interaction models allow us to quantify differences in AA or DD between *responsive* and *non-responsive* bats with increasing temperatures. The response variables (AA and DD) are evaluated in relation to temperature and response type as fixed factors, with individuals included as random effects. The *Response Type* factor comprises two categories: *responsive* or *non-responsive* bats. The results of this model could be interpreted as follows: 1) a significant effect of the factor *Response Type* would suggest that mean values of AA and/or DD differ between *responsive* or *non-responsive* bats. 2) A significant effect of *Temperature* would be interpreted as changing values of AA and/or DD across temperatures. 3) Finally, a significant effect of the interaction (*Response Type * Temperature*) on AA and/or DD would suggest that *responsive* and *non-responsive* bats respond differently to increasing temperatures.

## RESULTS

Call peak frequency did not change with increasing ambient temperatures in any of the species (p = 0.223-0.918; Fig. 2; Table 1). Likewise, apparent source level (apSL) did not change for most *Myotis* species, but it decreased for *M. oxyotus* (p=0.020) and for *E. brasiliensis* (p=0.043) with increasing temperatures (Fig. 2, Table 1). Call duration increased with temperature in *M. pilosatibialis* (p=0.002) and *M. oxyotus* (p=0.016); potentially also in *M. nigricans/elegans* (p=0.081), but not in the other two species, *E. brasiliensis* and *M. riparius* (Fig. 2, Table 1).

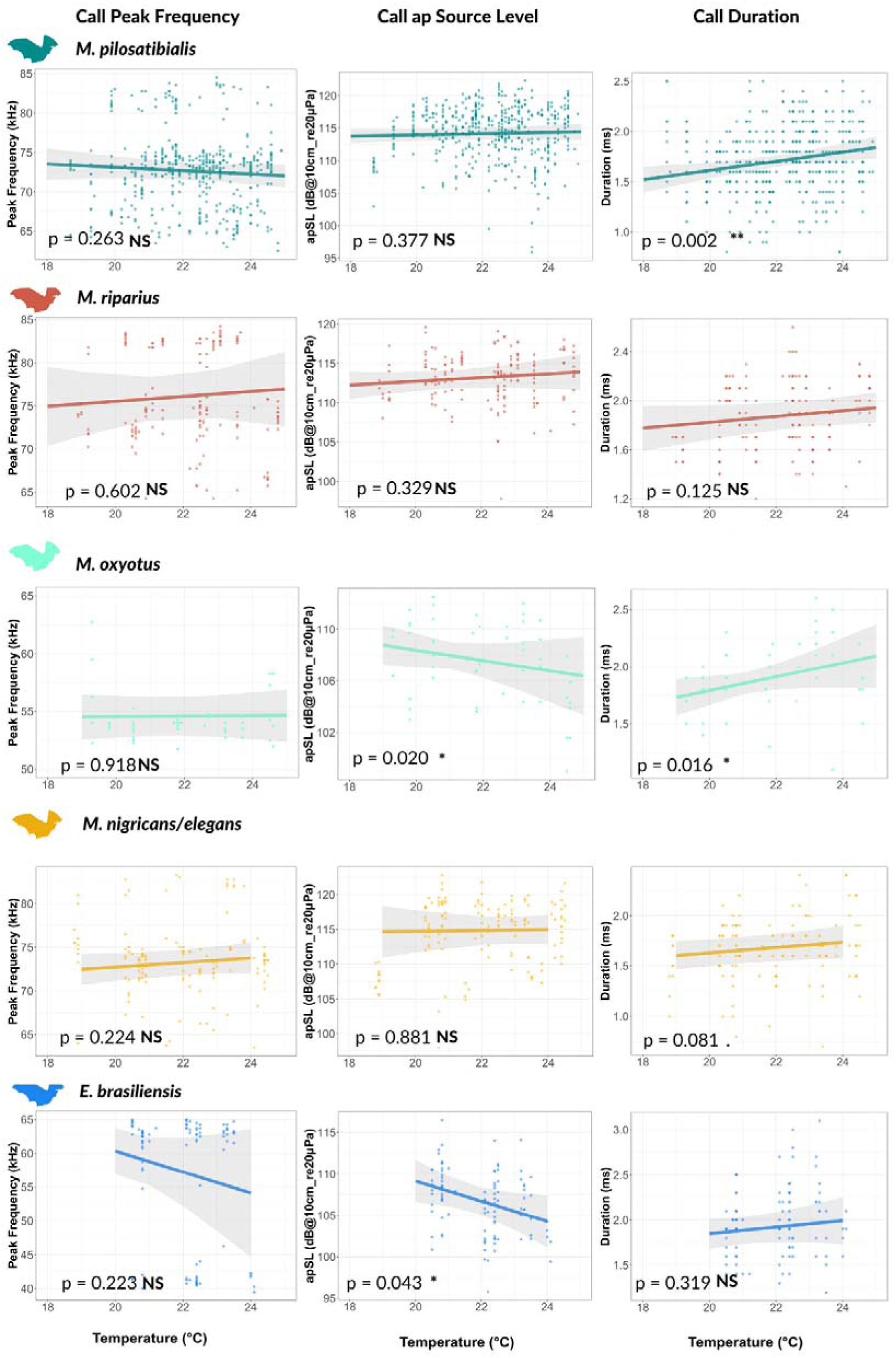

To quantify the combined effects of call frequency and weather conditions on atmospheric attenuation (AA), we compared the AA experienced by *non-responsive* bats with the AA experienced by *responsive* bats (Fig. 3, Table 2). *M. nigricans/elegans* (p=0.001) and *M. riparius* (p=0.026) experienced an increasing AA with increasing temperatures (Table 2), but not the other two *Myotis* species (p=0.321 for *M. oxyotus*; p=0.140 for *M. pilosatibialis)*, or *E. brasiliensis* experienced a decreasing AA with increasing temperatures (p=0.044, Table 2, Fig. 3). This pattern of AA as a function of temperature was the same for *responsive* and *non-responsive* bats for all *Myotis* species (Table 2, Fig. 3, interaction AA * Temperature, p = 0.178 – 0.816), matching the mostly lacking or only minor changes in call parameters. In contrast, in *E. brasiliensis*, the pattern of AA as a function of temperature differed between the *non-responsive* and *responsive* bat (Fig. 3, interaction AA * Temperature, p= 0.015): while AA decreased for the *responsive* bat from 2.2 to 1.9 dB/m with increasing temperatures, it was constant for the *non-responsive* bat at 2.2 dB/m across temperatures (Table 2). The decrease in AA for the *responsive* bat was likely driven by two factors: (1) the bat’s reduction of peak frequency from 60.3 kHz (at 20°) to 54.0 kHz (at 24°), albeit non-significant (Fig. 2) combined with (2) the effect of increasing temperatures which also causes a slight reduction of AA at this bats’ call frequencies of 55-60 kHz and the prevalent ambient weather conditions (20-24°C, >60% RH; Goerlitz, 2018).

**Figure 3:**
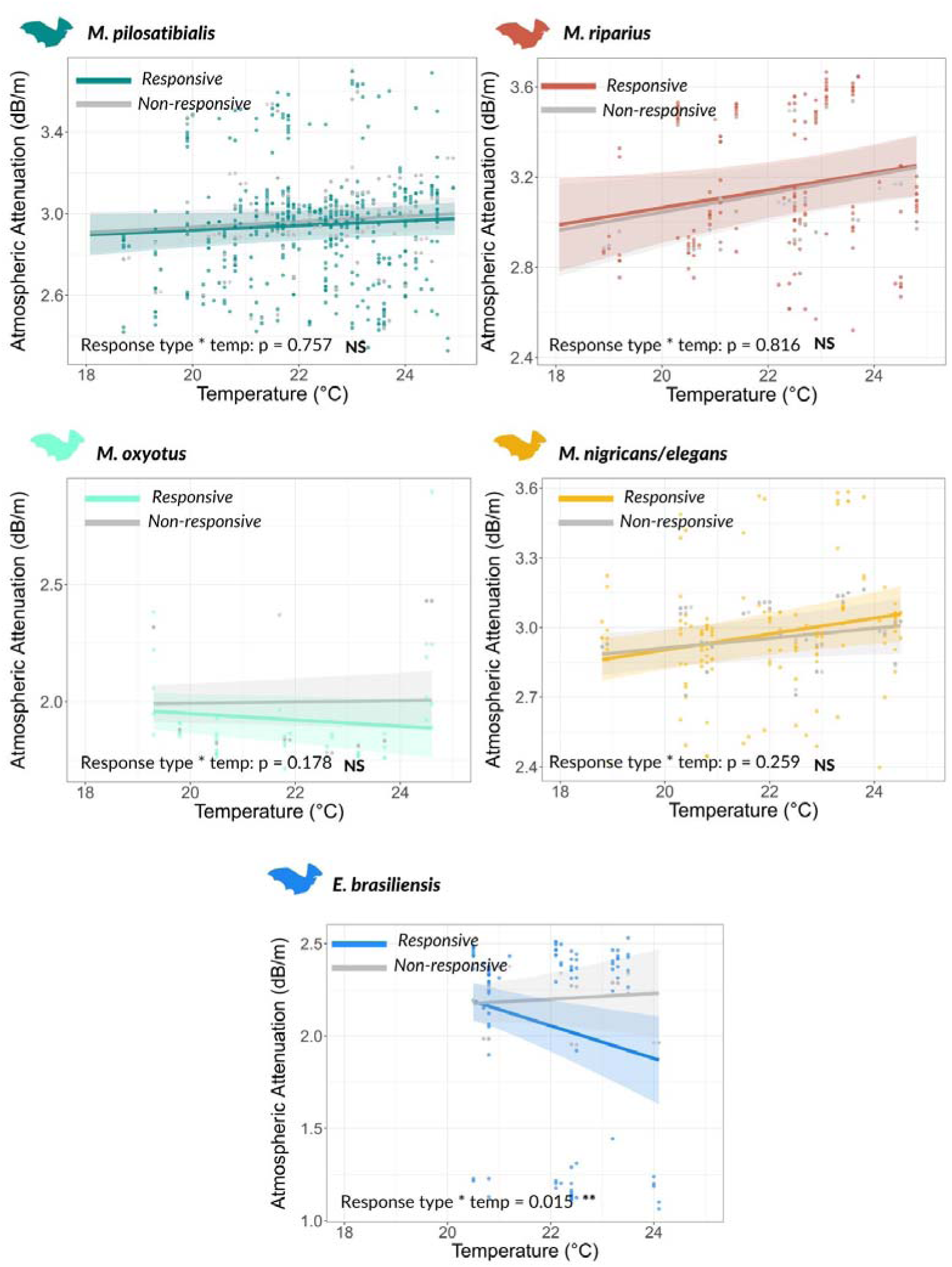
Atmospheric attenuation as a function of increasing temperatures, both for measured *responsive* (coloured) and hypothetical *non*-*responsive* (grey) bats. Dots show individual data, either calculated for actual call parameters and weather conditions (*responsive* bat: colored) or for call parameters at ambient temperature and actual weather data (*non-responsive* bats: grey). Lines are model results with the 95% confidence interval (shaded region). P-values in each panel indicate if the slope of AA over increasing temperature differs between *responsive* and *non-responsive* bats.

**Table 2.** Mean values and confidence intervals of atmospheric attenuation (AA) per experimental temperature. P-values for the interaction (int) show whether there are differences between *responsive* and *non-responsive* bats with changes in temperature. Arrows represent the difference between values of *responsive* and *non-responsive* bats across temperatures, with blue representing no difference, and red indicating that AA is lower in *responsive* bats. The p-values for temperature (temp) show whether AA changes with temperatures, with arrows representing an increase (green), no change (blue), and a decrease (red).

|  |  | Atmospheric Attenuation (-dB/m) |  |  |  |
| --- | --- | --- | --- | --- | --- |
|  |  | Mean (95% CI) |  | p-value |  |
| Species |  | *Responsive* | *Non-responsive* | int | temp |
| *M. pilosatibialis* | Ta°C | 2.9 (2.8, 3.0) | 2.9 (2.8, 3.0) | 0.757 | 0.140 |
|  | +2 °C | 2.9 (2.9, 3.0) | 3.0 (2.9, 3.0) | → | → |
|  | +4C | 3.0 (2.9, 3.1) | 3.0 (2.9, 3.1) |  |  |
| *M. riparius* | Ta°C | 3.0 (2.8, 3.0) | 2.9 (2.7, 3.2) | 0.816 | 0.026 |
|  | +2 °C | 3.1 (3.0, 3.2) | 3.1 (2.9, 3.2) | → | ↑ |
|  | +4C | 3.3 (3.1, 3.4) | 3.3 (3.1, 3.4) |  |  |
| *M. oxyotus* | Ta°C | 2.0 (1.8, 2.1) | 2.0 (1.8, 2.2) | 0.178 | 0.321 |
|  | +2 °C | 1.9 (1.7, 2.1) | 2.0 (1.8, 2.2) | → | → |
|  | +4C | 1.9 (1.6, 2.2) | 2.0(1.7, 2.3) |  |  |
| *M. nigricans/elegans* | Ta°C | 2.8 (2.8, 2.9) | 2.9 (2.8, 3.0) | 0.259 | 0.001 |
|  | +2 °C | 3.0 (2.9, 3.0) | 2.9 (2.9, 3.0) | → | ↑ |
|  | +4C | 3.1 (2.9, 3.2) | 3.0 (2.9, 3.1) |  |  |
| *E. brasiliensis* | Ta°C | 2.2 (2.0, 2.3) | 2.2 (2.0, 2.3) | 0.015 | 0.044 |
|  | +2 °C | 2.0 (1.8, 2.3) | 2.2 (2.0, 2.4) | ↓ | ↓ |
|  | +4C | 1.9 (1.5, 2.2) | 2.2 (1.9, 2.6) |  |  |

To quantify the contribution of the combined changes of call parameters and weather conditions on prey detection distance (DD), we compared the DD for prey by *non-responsive* bats with that of *responsive* bats (Fig. 4, Table 3). DD was constant across temperatures for most *Myotis* species, both *responsive* and *non-responsive* (Fig. 4, Table 2), except for *M. riparius*, where DD increased from 1.29 to 1.36 m in *responsive* bats (Table 3, Fig. 4). Since detection distance is influenced both directly by the call’s apparent source level, and to a lesser degree by its duration, and indirectly by call frequency through its effect on AA, the similarity between *responsive* and *non-responsive* values is expected given the mostly absent effects of temperature on call parameters (Fig. 2) and AA (Fig. 3). In the case of *M. riparius*, the differences in the slopes of DD across temperatures (p = 0.055) resulted in a slight increase in detection distance for the *responsive* bat with increasing temperatures. Finally, and as for AA, the slopes of DD over temperature differed between the *responsive* and *non-responsive E. brasiliensis* bat (p = 0.008). For the *responsive* bat, DD decreased with increasing temperatures from 1.22 to 1.05 m (Fig. 4 and Table 3). These results on DD were calculated for a medium sized prey (20mm^2^). Increasing or decreasing prey size by 10 mm^2^, resulted in a corresponding in- or decrease of DD by ∼0.5 m (Table S2).

**Figure 4:**
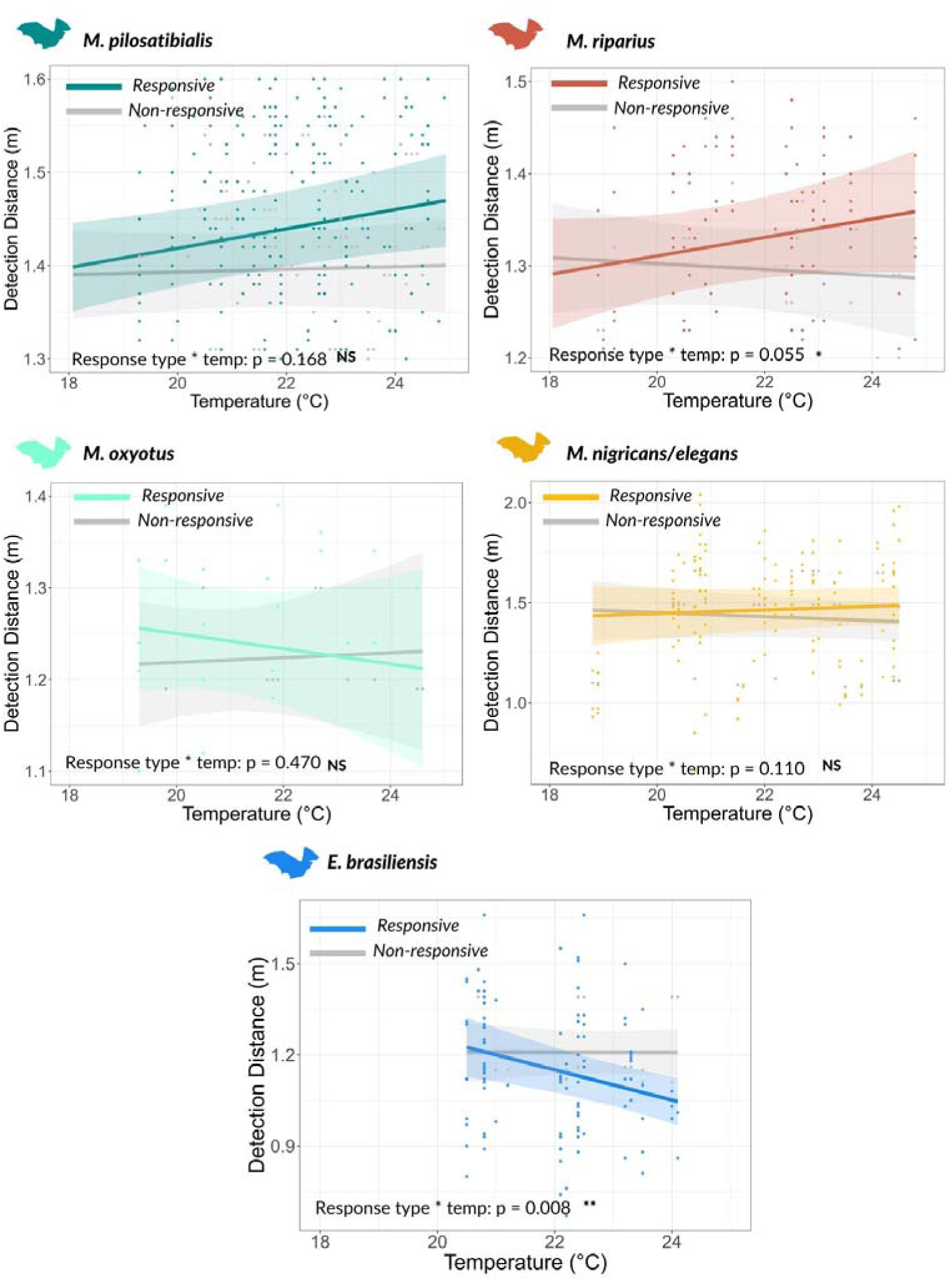
Detection distance as a function of increasing temperature, both for measured responsive (coloured) and hypothetical *non-responsive* (grey) bats. Dots show individual data, either calculated for actual call parameters and weather conditions (*responsive* bat: coloured) or for call parameters at ambient temperature and actual weather data (*non-responsive* bats: grey). Lines are model results with 95% confidence interval (shaded region). P-values in each panel indicate if slope of DD over increasing temperature differs between *responsive* and *non-responsive* bats.

**Table 3.**
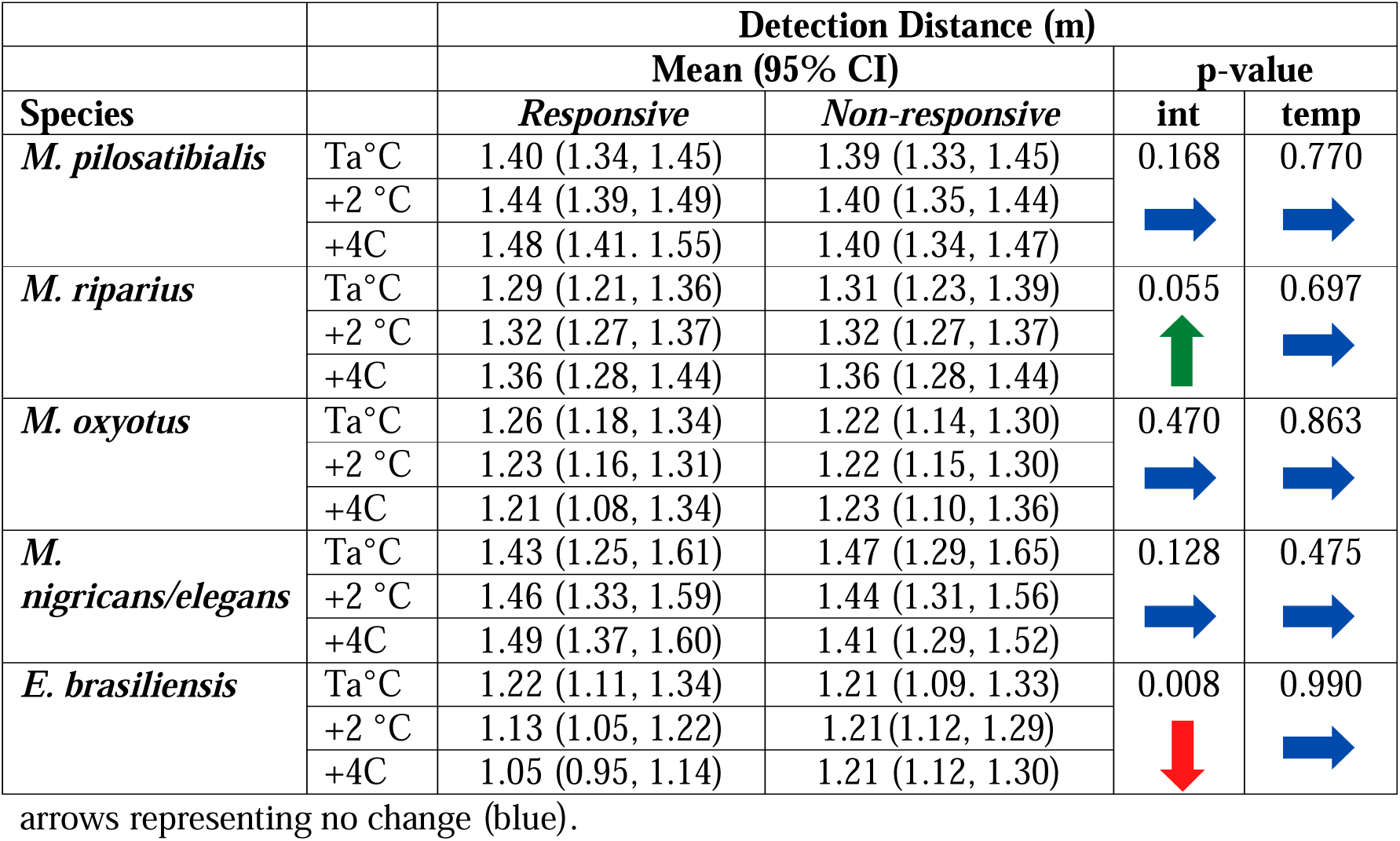
Mean values and confidence intervals of detection distance (DD) per experimental temperatures. P-values for the interaction (int) show whether there are differences between *responsive* and *non-responsive* bats with changes in temperature. Arrows represent the difference between values of *responsive* and *non-responsive* bats across temperatures, with green indicating that DD is larger in *responsive* bats, blue representing no difference, and red indicating that DD is lower in *responsive* bats. The p-values for temperature (temp) show whether DD changes with temperatures, with

|  |  | Detection Distance (m) |  |  |  |
| --- | --- | --- | --- | --- | --- |
|  |  | Mean (95% CI) |  | p-value |  |
| Species |  | *Responsive* | *Non-responsive* | int | temp |
| *M. pilosatibialis* | Ta°C | 1.40 (1.34, 1.45) | 1.39 (1.33, 1.45) | 0.168 | 0.770 |
|  | +2 °C | 1.44 (1.39, 1.49) | 1.40 (1.35, 1.44) |  |  |
|  | +4C | 1.48 (1.41, 1.55) | 1.40 (1.34, 1.47) |  |  |
| *M. riparius* | Ta°C | 1.29 (1.21, 1.36) | 1.31 (1.23, 1.39) | 0.055 | 0.697 |
|  | +2 °C | 1.32 (1.27, 1.37) | 1.32 (1.27, 1.37) |  |  |
|  | +4C | 1.36 (1.28, 1.44) | 1.36 (1.28, 1.44) |  |  |
| *M. oxyotus* | Ta°C | 1.26 (1.18, 1.34) | 1.22 (1.14, 1.30) | 0.470 | 0.863 |
|  | +2 °C | 1.23 (1.16, 1.31) | 1.22 (1.15, 1.30) |  |  |
|  | +4C | 1.21 (1.08, 1.34) | 1.23 (1.10, 1.36) |  |  |
| *M. nigricans/elegans* | Ta°C | 1.43 (1.25, 1.61) | 1.47 (1.29, 1.65) | 0.128 | 0.475 |
|  | +2 °C | 1.46 (1.33, 1.59) | 1.44 (1.31, 1.56) |  |  |
|  | +4C | 1.49 (1.37, 1.60) | 1.41 (1.29, 1.52) |  |  |
| *E. brasiliensis* | Ta°C | 1.22 (1.11, 1.34) | 1.21 (1.09, 1.33) | 0.008 | 0.990 |
|  | +2 °C | 1.13 (1.05, 1.22) | 1.21(1.12, 1.29) |  |  |
|  | +4C | 1.05 (0.95, 1.14) | 1.21 (1.12, 1.30) |  |  |

arrows representing no change (blue).

## DISCUSSION

Echolocation is highly dynamic, and bats adapt their calls constantly to changing conditions. Thus, we expected adaptive changes in the echolocation calls from our studied species to counteract the predicted reduction of prey detection distance caused by increasing atmospheric attenuation (AA) because of changing ambient temperatures. As predicted and published before (Barclay et al., 1999; Russo et al., 2018; Wund, 2006), we found inter-individual variation in call parameters within and between the different experimental temperatures (Figs. S1-S3). Most notably, we found a consistent and significant increase in call duration in two of five species. In contrast, the predicted reduction in call peak frequency (PF) and increase in apparent source level (aSL) were generally absent, or even opposed, for example by a decrease in aSL in *M. oxyotus* and *E. brasiliensis*.

Our experimental increase of ambient temperatures only caused a significant increase of AA in two species, *M. riparius* and *M. nigricans/elegans* (Table 2); which, however, did not prompt the expected decrease in call peak frequency and/or an increase in source level. We also found that detection distance remained relatively constant with increasing temperature. These results could provide further evidence that alterations in detection distance, whether increasing or decreasing, may be less pronounced in tropical species compared to temperate ones (Luo et al., 2014). However, additional studies are necessary to better understand the effect of rising temperatures on detection distance for a larger number of species and call types within species.

One reason for this lack of change in AA in our study might be the relatively low variation in temperature. For example, in a temperate habitat with strong variations in temperature and humidity (> Δ16°C and > Δ40% RH, respectively), AA increased by 0.7 dB/m (de Framond, et al., 2023). In contrast, in our study, variation in weather conditions was considerably smaller (< Δ7°C and < Δ25% RH), causing AA to change by only 0.1-0.3 dB/m, resulting in a two-way echo-level reduction of only 0.3-0.6 dB over the modelled prey detection distance of ∼1.5 m. This appears too small to have a significant effect on prey detection. Our experimental increase of temperature by 2 and 4°C thus had likely no relevant effect on AA and prey detection distances. This suggests in turn that anticipated increases in average temperature caused by climate change will likely not affect the sensory range of the bat species studied here, at least under the prevailing relative humidity levels at our study site.

Relative humidity also influences AA and thus the maximum detection distances by bat echolocation (Goerlitz, 2018). Tropical forests in Central America are forecasted to suffer a reduction in precipitation as a result of climate change (Lyra et al., 2017), with significant drying trends in southern Costa Rica (Hidalgo et al., 2017). At the prevailing temperature conditions of our study site, increasing ambient temperatures will have a stronger increasing effect on AA when relative humidity is lower and for call frequencies around 55-75 kHz of the bats studied here (Goerlitz, 2018). To investigate the effect of lower relative humidity at the same increasing temperature values used in our experiments, we modelled changes in AA for *non-responsive* bats in each studied species, and we fixed relative humidity at 50%, 75% and 100%. We found for all species significantly divergent AA values (p<0.001) between hypothetical scenarios of low and medium relative humidity in comparison to a high relative humidity setting (Fig. S4). At 50% relative humidity, AA increased in all *Myotis* species by 0.6 dB/m with increasing temperatures; and increases by 0.3 dB/m at medium relative humidity (75%). It stayed constant at high relative humidity (100%, Table S1). This result matches the predictions of other theoretical and empirical studies (Goerlitz, 2018; Lawrence & Simmons, 1982; Snell-Rood, 2012), suggesting that Neotropical montane bat species with high frequency echolocation calls (<70 kHz) might suffer from reduced prey detection ability as mountain ecosystems become drier and warmer. Further studies will need to address whether bats experience stronger changes of AA under these conditions, and whether they will behaviorally adjust calls in response to drier conditions.

We expected bats to increase their call source level in response to increasing atmospheric attenuation. Only *E. brasiliensis* and *M. oxyotus* adjusted this parameter, yet lowered it with increasing temperature. While echolocation during flight poses no additional energetic costs (Speakman & Racey, 1991; Voigt & Lewanzik, 2012), this might not be the case for very high-intensity calls (> ∼110 dB SPL @ 10cm; Currie et al., 2020). Hence, bats may experience physiological constraints to increase call levels beyond a certain threshold. Similarly, bats may experience trade-offs when (strongly) reducing call frequency. By lowering call frequency, bats can increase detection distance, but in parallel confine prey-detection to larger items because smaller objects reflect lower frequencies less (de Framond, Reininger, et al., 2023; Jung et al., 2014). Excluding a part of the potential prey spectrum by reducing call frequency could reduce a bat’s foraging success. Lowering call frequency to maintain prey detection distances while excluding smaller prey items is a trade-off that is most likely context specific. For example, individuals that normally consume larger prey items would not suffer major losses when decreasing call frequency, whereas those that typically consume smaller prey would probably not be able to trade-off a large portion of available prey for an increase in detection distance. This might explain some of the variation observed in bats’ vocal responses in our study.

While most studies to date have investigated how bats may adapt frequency and source level to deal with changes in AA and other auditory challenges (de Framond, Reininger, et al., 2023; Snell-Rood, 2012; Surlykke & Kalko, 2008), only few have considered the effect of changing call duration on sound perception and detection distance (but see Chaverri & Quirós, 2017; Luo et al., 2014; Schmidt & Thaller, 1994). Our study provides additional support that adjusting call duration is another potential mechanism to improve signal detection. Increasing call duration improves signal detectability by about 6 dB per doubling of duration for short calls (Luo et al., 2014). Bats may increase call duration in noisy environments (Corcoran & Moss, 2017; Luo et al., 2015; Tressler & Smotherman, 2009), and our results suggest that bats may use the same mechanism to counteract reduced echo levels to improve detection distance. However, bats increased call duration by ∼0.1-0.4 ms for average call durations of ∼1.5-2.0 ms, i.e., by a factor of ∼1.05 - 1.27-fold, resulting in an increase in signal detectability of 0.2-0.9 dB. Given these small effect sizes, more studies will be needed to evaluate the relevance of call duration for improving signal detectability, and its dependence on other constraints, for example if changing frequency and source level might have interacting effects.

To our knowledge, this is the first experimental assessment of short-term adjustments of echolocation calls to experimentally raised ambient temperature in the Neotropical region, providing first data about a scarcely studied topic (Festa et al., 2022). Our results suggest that the average effect of warming on detection distance seems to be small for close-range prey detection, likely precluding the need for call adjustments in some bat species and under specific weather conditions. Nevertheless, future studies are needed to understand how call types, call function, behavioral context and ecology interact and affect sound perception in a wider range of species and weather conditions, and how bats deal with changes potentially challenging their perception. For example, in response to changing weather conditions, two species of molossid bats did not change their frequency-modulated calls that are used for close range object detection, and which are similar to the calls of the species in our study. In contrast, they adjusted their lower-frequency, constant-frequency calls that are used for long-range object detection (Chaverri & Quirós, 2017). Further research is needed to understand if other bat species will be affected by changing weather and climatic conditions and if they will be capable of adjusting their echolocation calls, their most important sensory input.

## Acknowledgements

We thank Silvia Chaves-Ramírez, Christian Castillo Salazar, Federico Granados, and Nazaret Rojas for their vital support during field work. We also thank to the Acoustic and Functional Ecology team at the Max Plank Institute, Seewiessen, Germany for their support with methodological ideas for this research, especially to Theresa Hügel, Thejasvi Beleyur and Daniel Lewansik. Especial thanks to Lena de Framond for their support and bold ideas during data analysis. Finally, we thank Las Cruces Biological Station and their staff for supporting this research.

## Author contributions

Conceptualization: P.I-P, G.C.; Methodology: P.I-P, H.R.G, G. C.; Formal analysis: P. I-P, G. C., M. A-S.; Investigation: P. I-P, G. C., H.R.G.; Resources: P. I-P, G.C.; Data Curation: P. I-P.; Writing – original draft: P. I-P; Writing – review & editing: G.C, H.R.G.; Supervision: G.C., H.R.G.; Project administration: P. I-P.

## Data availability

Data supporting this article are available from: https://github.com/morceglo/Neotropical-bats-climate-change-call-emission

